# Indirect Readout of DNA Controls Filamentation and Activation of a Sequence-Specific Endonuclease

**DOI:** 10.1101/585943

**Authors:** Smarajit Polley, Dmitry Lyumkis, N. C. Horton

## Abstract

Filament or run-on oligomer formation by enzymes is increasingly recognized as an important phenomenon with potentially unique regulatory properties and biological roles. SgrAI is an allosterically regulated type II restriction endonuclease that forms run-on oligomeric (ROO) filaments with enhanced DNA cleavage activity and altered sequence specificity. Here, we present the 3.5 Å cryo-electron microscopy structure of the ROO filament of SgrAI bound to a mimic of cleaved primary site DNA and Mg^2+^. Large conformational changes stabilize a second metal ion cofactor binding site within the catalytic pocket and facilitate assembling a higher-order enzyme form that is competent for rapid DNA cleavage. The structural changes illuminate the mechanistic origin of hyper-accelerated DNA cleavage activity within the filamentous SgrAI form. An analysis of the protein-DNA interface and the stacking of individual nucleotides reveals how indirect DNA readout within filamentous SgrAI enables recognition of substantially more nucleotide sequences than its low-activity form, thereby expanding DNA sequence specificity. Together, substrate DNA binding, indirect readout, and filamentation simultaneously enhance SgrAI’s catalytic activity and modulate substrate preference. This unusual enzyme mechanism may have evolved to perform the specialized functions of bacterial innate immunity in rapid defense against invading phage DNA without causing damage to the host DNA.

## INTRODUCTION

Filament formation by non-cytoskeletal enzymes is a newly appreciated phenomenon. Although first shown *in vitro* for the metabolic enzymes acetyl-CoA carboxylase and phosphofructokinase in the 1970s^1–3^, it was not until 40 years later that filament formation was demonstrated to modulate enzyme activity under physiological conditions^4,5^. At around the same time, filament formation was surprisingly discovered for the unfolded protein response RNase/kinase Ire1^6^ and for numerous other enzymes previously unknown to form such assemblies^7–9^. Independently, filament formation was proposed to explain the unusual biophysical and biochemical activities seen in the type II restriction endonuclease SgrAI^10^.

SgrAI is a sequence-specific type II DNA restriction endonuclease that is allosterically activated by the same substrate DNA. A primary recognition sequence (CR|CCGGYG, where R=A or G and Y=C or T, | denotes cleavage site) activates the enzyme by over 200-fold, which in turn also relaxes the enzyme’s DNA sequence specificity, resulting in the cleavage of an additional 14 secondary sequences (C<u>C</u>CCGGYG or <u>X</u>RCCGGY G, where X=A,G, or T) up to 1000-fold faster^10–14^. In the absence of DNA, SgrAI is a homodimer composed of two 37 kDa chains, each with a single active site^15–18^. The enzyme binds primary or secondary site DNA in 1:1 ratios, and the ensuing complex is referred to as a DNA-bound dimer (DBD). Since the primary site DNA acts both as an allosteric effector *as well as* a substrate for enzymatic cleavage, it is expected that activated SgrAI possesses at least two DNA binding sites, one for enzymatic cleavage and one for the allosteric effector. Early studies suggested the formation of an activated SgrAI/DNA complex composed of at least two DBDs^16,17^, thereby providing a functional complex with two DNA binding sites. However, assemblies containing many more DBDs were shown by analytical ultracentrifugation and ionmobility mass spectrometry^10,19^. Negative stain EM revealed filaments of varied lengths with left-handed helical symmetry that we call run-on oligomers (ROO)^20^.

The ROO filament structure was previously resolved to ~nanometer resolution by cryo-EM and helical reconstruction, that revealed left-handed helical symmetry with approximately 4 DBDs per turn^20^. This structure inspired a low-resolution mechanistic model for the enzymatic behavior of SgrAI, wherein binding to primary site DNA induces a conformational change that favors ROO filament formation, which in turn stabilizes the activated enzyme state capable of accelerated DNA cleavage (**Fig. 1A**). SgrAI DBD bound to a secondary site DNA disfavors the activated conformation, and hence ROO filament formation by SgrAI bound to secondary site does not appreciably occur (**Fig. 1B**). This explains why cleavage of secondary site sequences is negligible unless a primary site is present – in sufficient concentration or on the same contiguous DNA – to promote ROO filament assembly^10,12,14^. Filaments formed by SgrAI bound to primary site DNA will drive ROO assembly, incorporating DBD-containing secondary sites, activating SgrAI for DNA cleavage on both primary and secondary sites, and thereby expanding the enzyme’s DNA sequence specificity (**Fig. 1C**).

**Figure 1.**
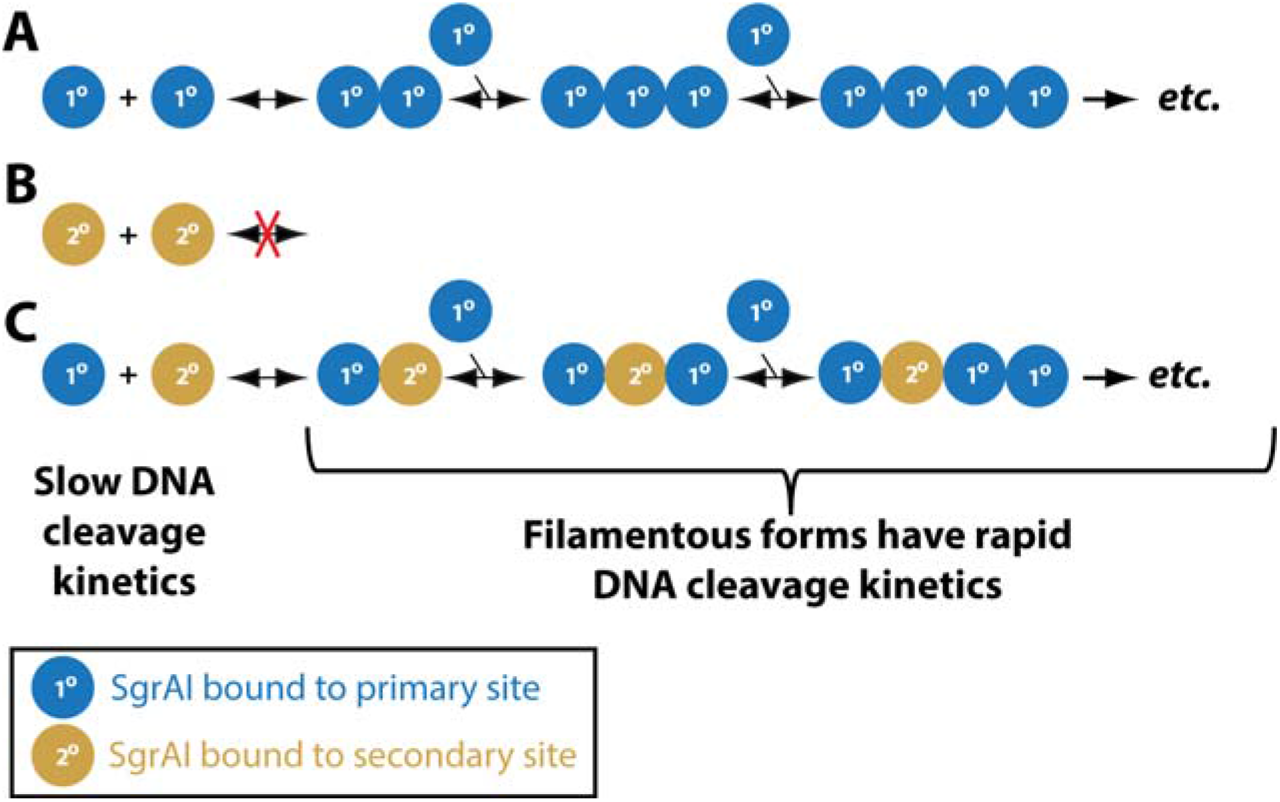
Schematic of differential behavior of SgrAI with primary and secondary site sequences. **A**. SgrAI bound to primary site DNA (cleaved or uncleaved, blue spheres) form ROO filaments with rapid DNA cleavage kinetics. The unfilamentous form cleaves DNA slowly. **B**. SgrAI bound to secondary site DNA only (gold) does not form filaments. **C**. SgrAI bound to secondary site DNA will form ROO filaments with SgrAI bound to primary site DNA, resulting in rapid DNA cleavage of both primary and secondary site sequences.

A full kinetic analysis of SgrAI-mediated cleavage of primary site DNA enabled the estimation of individual rate constants for the major steps in the reaction pathway, such as DBD association into the ROO filament, cleavage of DNA within the filament, DBD dissociation from the ROO filament, and dissociation of cleaved DNA from SgrAI^21,22^. These studies show that assembly of DBDs into the ROO filament is rate limiting when recognition sites are on separate DNA molecules, but fast when on the same contiguous DNA. Kinetic simulations show that this rate limiting step underlies the enzyme’s ability to sequester potentially damaging secondary site cleavage events onto invading phage DNA and away from the host, leaving host sites largely untouched^23^. The study also shows that DNA cleavage is fast, followed by a relatively slower disassembly of DBDs from the ROO filament. Although slower than DNA cleavage, the rate of disassembly is still sufficiently fast to prevent trapping of cleaved DNA within the filament. Furthermore, simulations also show that the very nature of filament formation imparts beneficial properties to the enzyme, both in terms of faster enzyme kinetics and its ability to sequester DNA cleavage onto phage and away from host, providing protection against infection and host DNA damage, respectively^23^. Importantly, mutations disrupting the ROO filament (even moderately) abolish all protection provided by SgrAI against phage infection, indicating that the rapid speed provided by ROO filament formation is critical to anti-phage activity^23^.

The previous low-resolution cryo-EM structure shows the general architecture of the ROO filament, but left open two important questions: 1) what is the mechanism for activated DNA cleavage? and 2) how do primary vs. secondary site DNA sequences differentially stabilize the activated conformation of SgrAI and the ROO filament? To address these questions and clarify the molecular mechanisms of enzyme activation and allosteric regulation, we determined the structure of activated, filamentous SgrAI to near-atomic resolution. The 3.5 Å cryo-EM maps allowed detailed model building and refinement, and enabled us to compare the ROO filament structure with previously determined x-ray crystal structures of the inactive, non-filamentous forms. The data revealed how numerous conformational changes in protein and DNA stabilize the SgrAI ROO, activate DNA cleavage, and how a single base pair change in secondary sites may modulate enzyme activation.

## RESULTS AND DISCUSSION

### Cryo-EM helical analysis of SgrAI ROO filaments resolves the activated enzyme form

To elucidate the molecular determinants of SgrAI activation, we sought to resolve the ROO filament to near-atomic resolution. We prepared ROO filaments from purified, recombinant, his-tagged wild type SgrAI^14^ bound to a 40-bp oligonucleotide DNA containing a pre-cleaved primary site sequence (PC DNA, see Methods). Cryo-EM micrographs revealed a large variety of differently-sized filaments, as expected for a ROO (**Fig. S1A**). Because the filaments were typically limited to ~<10 DBDs in size, thereby making it challenging to precisely define their orientations on the cryo-EM micrographs through conventional manual filament tracing, we sought to process the data in a single-particle manner. Template-based particle detection using a small distance between neighboring picks allowed us to select virtually all filamentous regions on the micrographs. 2D and 3D classification – without imposing helical symmetry – allowed us to separate the particle picks into compositionally distinct classes (**Fig. S1B-C**). Subsequent imposition of helical symmetry facilitated obtaining a high-resolution cryo-EM electrostatic potential map, resolved globally to 3.5 Å, which in turn enabled deriving an atomic model (**Fig. 2A, Fig. S1D-H**). The model is consistent with the final cryo-EM map, and has good geometry and statistics (**Fig. S1F, Fig. S2, and Table S1**).

**Figure 2.**
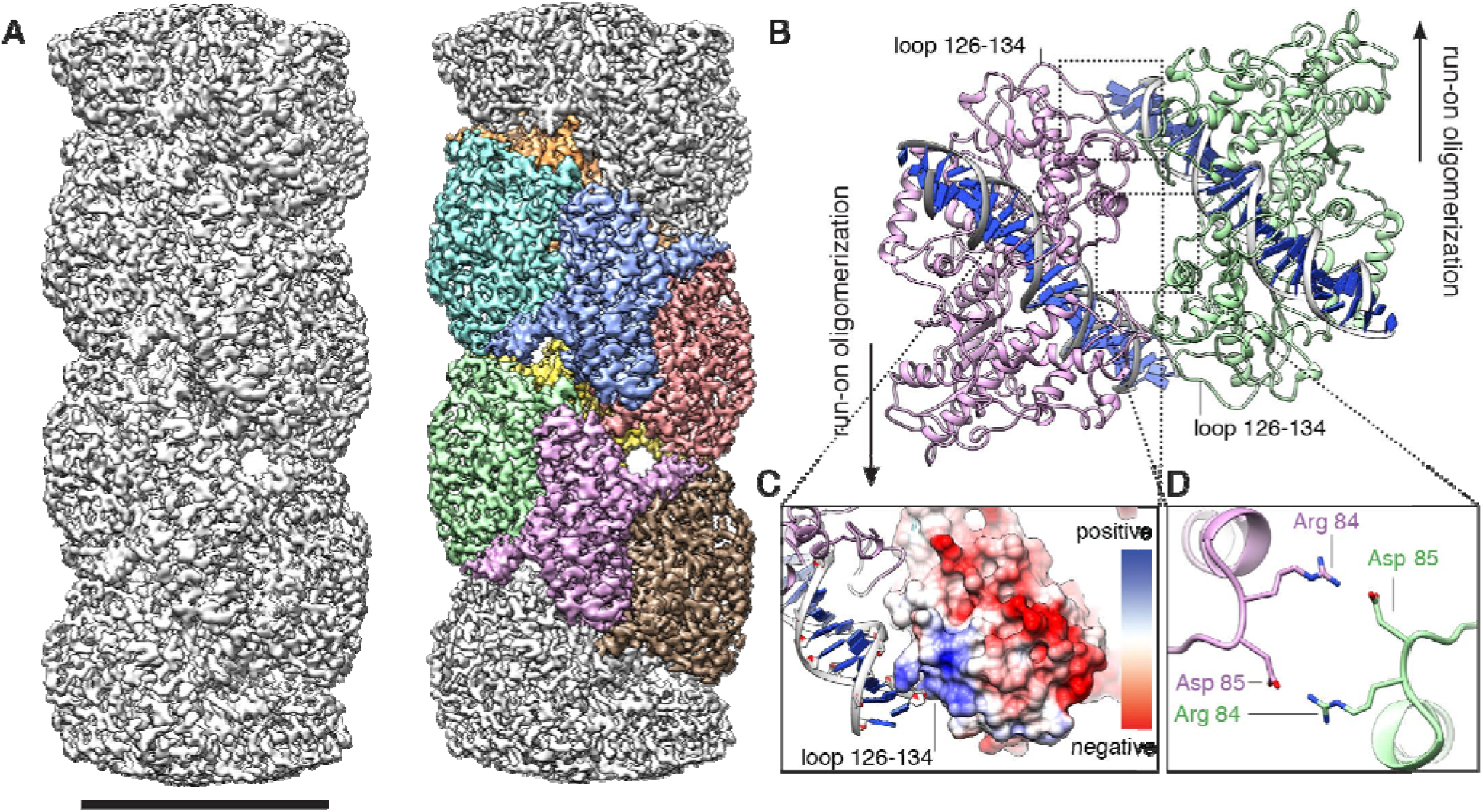
Structure of activated and oligomeric SgrAI. **(A)** Cryo-EM structure of activated SgrAI ROO, reconstructed to 3.5 Å with application of helical symmetry. At right, individual DBDs are colored onto the structure. Scale bar is 100 Å. **(B)** Atomic model of two DBDs, as viewed from the center of the helical axis. The filament can oligomerize in two directions. **(C-D)** Close-up views of interfaces forming inter-DBD contacts are indicated, including **(C)** electrostatic interactions between loop ~126-134 with flanking DNA and **(D)** salt-bridges formed by charged side-chains.

Each DBD constitutes the basic building block of activated and filamentous SgrAI. The enzyme can oligomerize in a run-on manner, from either side along the helical axis (**Fig. 2B**). In previous experiments, this property has been shown to specifically benefit the enzyme’s ability to quickly sequester phage DNA within the oligomeric form for rapid cleavage^24^. Within the filament, SgrAI makes several inter-DBD contacts between asymmetric units. The loop spanning residues 126-134 interfaces with flanking DNA. Electrostatic interactions from charged amino acids within this patch of the protein help maintain the oligomeric enzyme form (**Fig. 2C**). This patch also makes up a portion of the allosteric interface of the enzyme. Immediately alongside, loop 55-62 also packs against the flanking DNA, although this interface is primarily maintained by weaker van der Waals interactions with the DNA backbone. Several regions of the NTD make protein-protein interactions between neighboring DBDs within the central filament axis, including clusters of salt bridges that help to facilitate helical packing. Arg84 from DBD_n_ makes a salt bridge with Asp85 from DBD_n+1_ (**Fig. 2D**). Several other charged residues, including Arg11 and Glu8, reside immediately nearby, although their exact contributions cannot be accurately determined at the current resolution. These charged regions from multiple neighboring DBDs, both immediately alongside and at the opposing side of the helix, generally contribute to interface packing along the central helical axis. These data suggest that the interactions made within the central filament axis are specific, and are designed to facilitate oligomerization. The structural organization, combined with previous biochemical and functional data, implies that filament formation is an inherent property of the enzyme.

### The activated DNA-bound SgrAI enzyme exhibits multiple conformational rearrangements that stabilize the filamentous ROO form

To determine the molecular changes underlying enzyme activation, we compared the structure of the high activity ROO filament to a previously published X-ray structure of a low-activity, non-filamentous SgrAI DBD (PDB code 3DVO^18^). Superposition of the two DBDs reveals a ~9° rotation of one chain relative to the other (**Fig. 3A and Movie S1**), with shifts also occurring in the dimeric interface to accommodate this rotation (**Fig. 3B**). Such conformational changes allow for favorable interactions to occur within the ROO filament. Specifically, conformational changes place residues 84-87 and 22-34 into position for filament formation (**Fig. 3C**). Without such structural changes, the subunits would clash sterically at the interface marked by a red “X” in **Fig. 3C**, and thus these residues must facilitate proper helical packing in the filament. Changes in conformation also occur in the bound DNA, and result in favorable interactions between the base pairs flanking the recognition sequence and SgrAI residues (56-57 and 127-134) of a neighboring DBD in the ROO filament. **Figure 3D** show these interactions, comparing the ROO filament structure (in magenta, neighboring SgrAI in green), the non-filamentous low-activity DBD (gray), and an idealized B-form DNA (tan). The ROO DNA is bent towards neighboring SgrAI in the ROO filament relative to the idealized B-form DNA, but takes on a different path compared to the DNA in the non-filamentous enzyme assembly. Furthermore, DNA bending allows for interactions with residues 56-57 and 127-134 of the neighboring SgrAI, without steric conflicts. These residues have been shown to be important for enzyme activation, presumably by stabilizing the ROO filament^14^. Overall, the structural rearrangements observed within the filamentous SgrAI ROO (Movie S1) prevent unfavorable clashes and promote interactions between neighboring DBDs.

**Figure 3.**
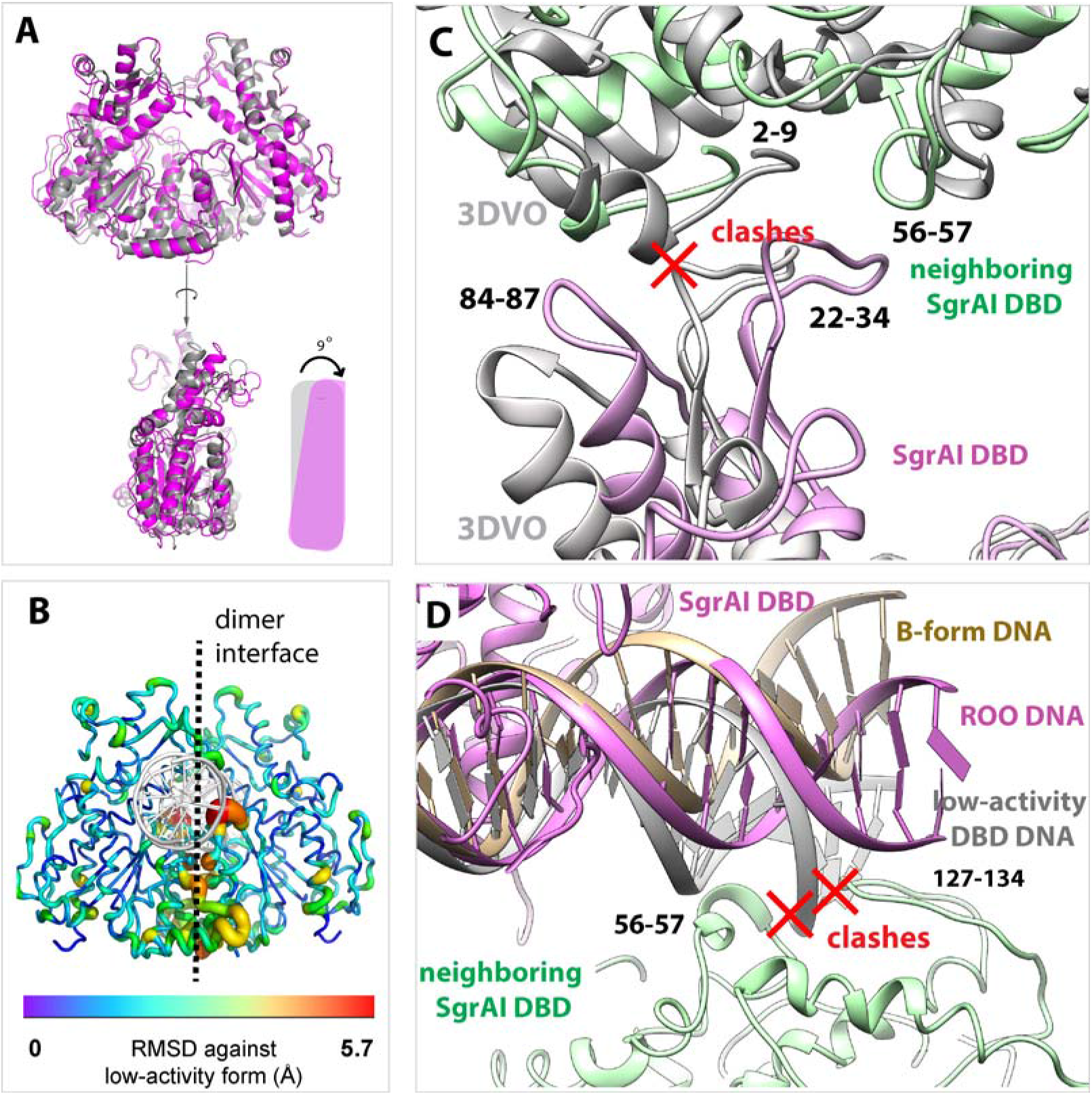
Conformational changes facilitate interactions in the ROO filament. **A**. Superposition of the low-activity DBD (3DVO, gray) and the cryo-EM (magenta) structures, with a schematic of the global conformation change. DNA not shown for clarity. **B**. Cartoon rendering of the SgrAI ROO DBD colored by RMSD against the low-activity form. **C**. Protein conformational changes in the ROO prevent steric clashes, which would occur in the low-activity form (marked by “X” in red). **D**. Comparison of flanking DNA between the ROO dimer, the low-activity X-ray structure extended using B-form DNA, and an idealized 40 bp B-form DNA (tan), extended from the 8 bp recognition sequence. In the cryo-EM structure, the DNA of one DBD takes on a distinct path within the filament in order to make contacts with neighboring SgrAI residues 127-134 and 56-57 (green) and prevent steric clashes.

### Mechanism of SgrAI hyper-activation for DNA cleavage through stabilization of a second Mg^2+^ binding site

Many nucleases, as well as other phosphoryl transfer enzymes, are dependent on divalent metal ion cofactors such as Mg^2+^ for their activity. SgrAI is no exception, and cleaves nucleic acids in the presence of Mg^2+^, and will also utilize Mn^2+^ or Co^2+^ for catalysis^18,25^. The now classical two-metal ion mechanism was first proposed for divalent ion-dependent DNA hydrolytic cleavage based on structures of a 3’-5’ DNA exonuclease^26^, and has since been used as a mechanistic model for many metal ion-dependent phosphoryl transfer enzymes^27–29^. Based on the activated cryo-EM reconstruction of filamentous SgrAI, the two-metal ion mechanism, adapted for the enzyme, is shown schematically in **Figure 4A** with experimental structural data supporting the mechanism in **Figure 4B**. In describing both panels, we have utilized the “tense” T and “relaxed” R states – terms adopted based on early enzymology work with hemoglobin^30^ – to describe the low- and high-activity forms of the enzyme, respectively.

**Figure 4.**
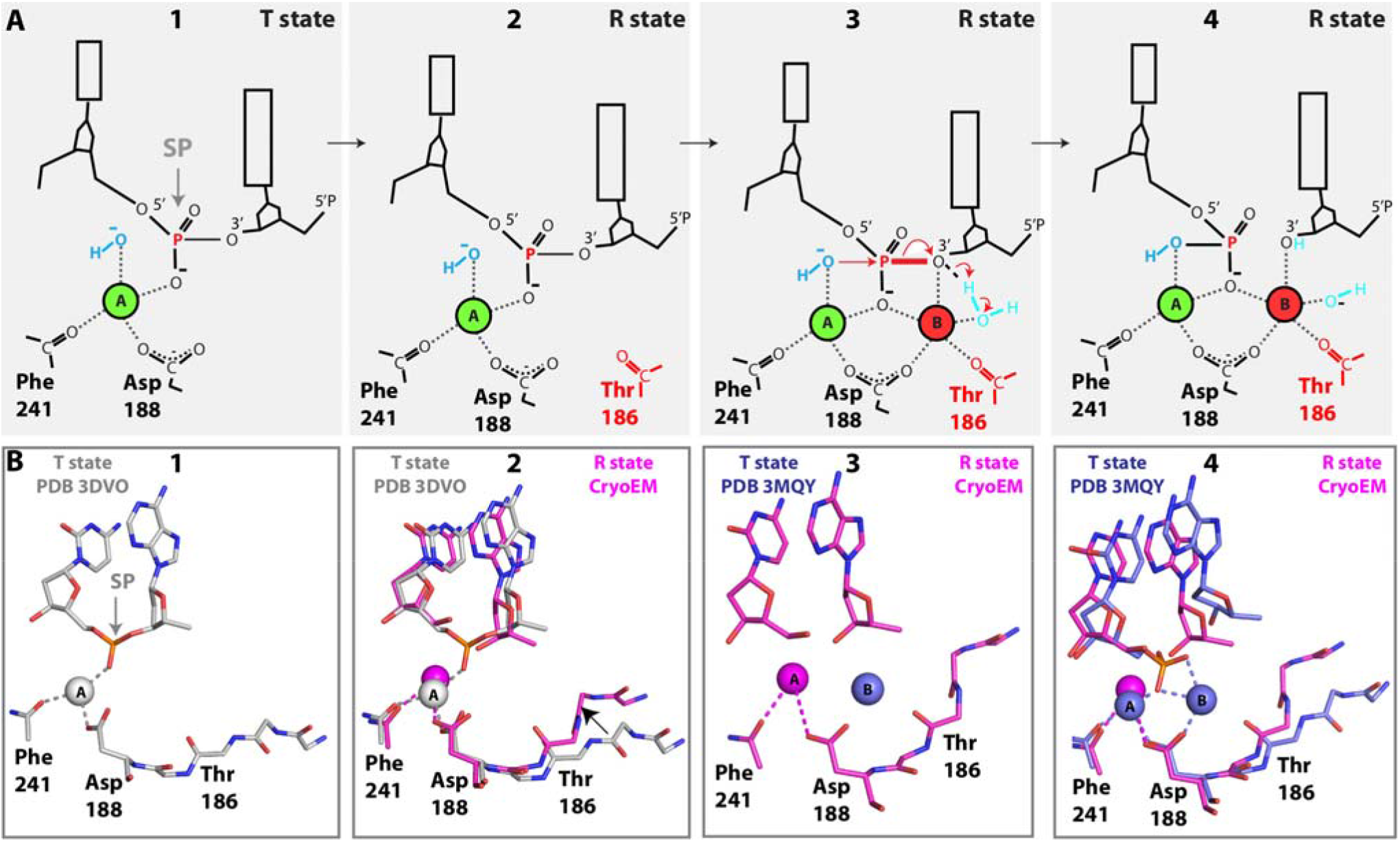
Active site configuration and two-metal ion cleavage mechanism. **A.** Schematic of the two-metal ion mechanism adapted for SgrAI. Panel 1 shows the low activity (T state) conformation where only metal ion Site A is occupied. Panel 2 depicts the R state conformation, based on the cryo-EM ROO filament structure, with the backbone carbonyl of Thr186 shifted closer to stabilize the metal ion of Site B. Panel 3 shows the R state with Site B occupied and just prior to the cleavage reaction. The scissile phosphodiester (SP) containing the bond to be cleaved (red thick line) is indicated. Panel 4 shows the product complex in the R state. **B**. Representative active site structures. Panel 1 shows the non-filamentous x-ray crystal structure of SgrAI bound to uncleaved primary site DNA and Ca^2+^ occupying Site A (PDB 3DVO), the T state. Panel 2 shows the active site in the cryo-EM ROO filament structure with Mg^2+^ occupying Site A in magenta (the R state), and 3DVO as shown in panel 1B (the T state). An arrow shows the shift in residues near Thr 186. Panel 3 shows the cryo-EM structure as in Panel 2 (R state, magenta) and the Site B Mg^2+^ (blue), identified in the post-catalytic crystal structure of SgrAI bound to cleaved primary site DNA, in the low activity T state conformation (PDB 3MQY). Panel 4 shows the complete active site arrangement of the post-catalytic product from the low activity T state overlaid on the cryo-EM ROO model. The superposition highlights the large distance spanned by the carbonyl of Thr186 and the Site B Mg^2+^ in the low-activity T state (PDB 3MQY), as compared to the high-activity R state (cryo-EM ROO).

In the first panel, SgrAI is in the low activity T state, and only metal ion Site A is occupied. This metal ion ligates both a nonesterified oxygen of the scissile phosphate (SP, **Fig. 4A**, panel 1), as well as a water molecule, inducing its deprotonation to hydroxide (blue). The previous X-ray crystal structure of SgrAI bound to uncleaved primary site DNA and Ca^2+^ (PDB file 3DVO^18^), shows this state (**Fig. 4B**, panel 1). The next panel of **Figure 4A**, panel 2, shows the shift in conformation to the activated R state, which brings the backbone carbonyl of Thr 186 closer to the metal ion binding Site B. The current cryo-EM ROO filament structure of SgrAI bound to Mg^2+^ and primary site DNA (but missing the SP) shows this state (**Fig. 4B**, panel 2). In this structure, the Site B Mg^2+^ is unoccupied, presumably due to the absence of the SP. The shift in residues 184-187 is shown by the black arrow. Panel 3 of **Figure 4A** shows the active site immediately prior to the reaction. Once Mg^2+^ binds Site B, it ligates the SP via the same nonesterified oxygen as the Site A Mg^2+^, as well as the leaving group, the O3’. Although currently there is no direct evidence for ligation between the Site B Mg^2+^ and the O3’, it is commonly proposed to occur in two-metal ion mechanisms of other enzymes^28^. The Site B metal ion also ligates a water molecule positioned to donate a proton to the O3’ leaving group (cyan, **Fig. 4A**, panel 3), thereby stabilizing the otherwise unfavorable negative charge that forms on the O3’ as the bond with phosphorus (red, **Fig. 4A**, panel 3) breaks. Panel 3 of **Figure 4B** shows the cryo-EM ROO filament structure in magenta, overlaid onto the Site B Mg^2+^ from a post-catalytic product structure of SgrAI bound to cleaved DNA and residing in the T state (PDB code 3MQY), in blue. The shift in residues 183-188 in the R state brings the backbone carbonyl of Thr 186 1.5 Å closer to the Site B metal ion (blue, **Fig. 4B**, panel 3), which would allow for stabilization of the ion either through direct or second shell ligation. Finally, panel 4 of **Figure 4A** shows the predicted product structure after the reaction has occurred. Panel 4 of **Figure 4B** shows the same product structure of 3MQY, with all regions displayed, superimposed on the cryo-EM ROO. In the T state conformation of 3MQY, the backbone carbonyl of Thr 186 is too far from the site B Mg^2+^ to directly coordinate the metal ion (**Fig. 4B**, panel 4). Presumably, this previously observed low-activity conformation included other structural differences in comparison to the expected enzyme configuration in the high-activity state. In contrast, the current cryo-EM reconstruction provides the first glimpse into the structural configuration of the enzyme active site in the hyper-activated R state that is competent for rapid DNA cleavage and an improved model for the post-catalytic product state of the hyper-activated form. Together with previous low-activity crystal structures, and building upon prior knowledge of the two-metal ion hydrolytic cleavage mechanism, these snapshots provide the most comprehensive insight into the full mechanism of catalytic cleavage activity for SgrAI.

### DNA distortions suggest that ROO-mediated sequence specificity expansion is influenced by indirect readout

To understand how DNA sequence affects SgrAI preferences for primary and secondary binding sites, we investigated the protein-DNA interactions and DNA structure in the inactive and active enzyme conformational states. The inactive state is represented by PDB 3DVO, and the active state is represented by the current cryo-EM ROO model. We could find no significant differences in direct readout or in other direct contacts to the 8-bp recognition sequence (**Fig. S3**). However, analysis of the DNA structure showed a large rearrangement at the center (4^th^) base step, between the C4 and G5 nucleotides in CAC<u>CG</u>GTG (**Fig. 5A**). The 4 Å rise at this base step is unusually high in both conformations (**Fig. 5A** right, the van der Waals rise in B form DNA is 3.4 Å). Such a large rise will likely weaken the stacking energy of these bases. Analysis of RMSD of the 8-bp recognition sequence between the two structures reveals better alignments when superimposing each 4-bp half-site, rather than the full 8-bp sequence (**Table S2**). Furthermore, calculation of the base overlap areas between neighboring dinucleotide pairs confirmed that the 4^th^ base step undergoes the largest structural change upon transitioning from the low to the high activity state (**Table S3**). Hence, it appears that the DNA accommodates the large conformational change in the SgrAI dimer, without disrupting important direct readout contacts between SgrAI and the DNA, by allowing each 4-bp half-site to move independently. The weakened stacking and large rise at the center base step allows for this independent movement.

**Figure 5.**
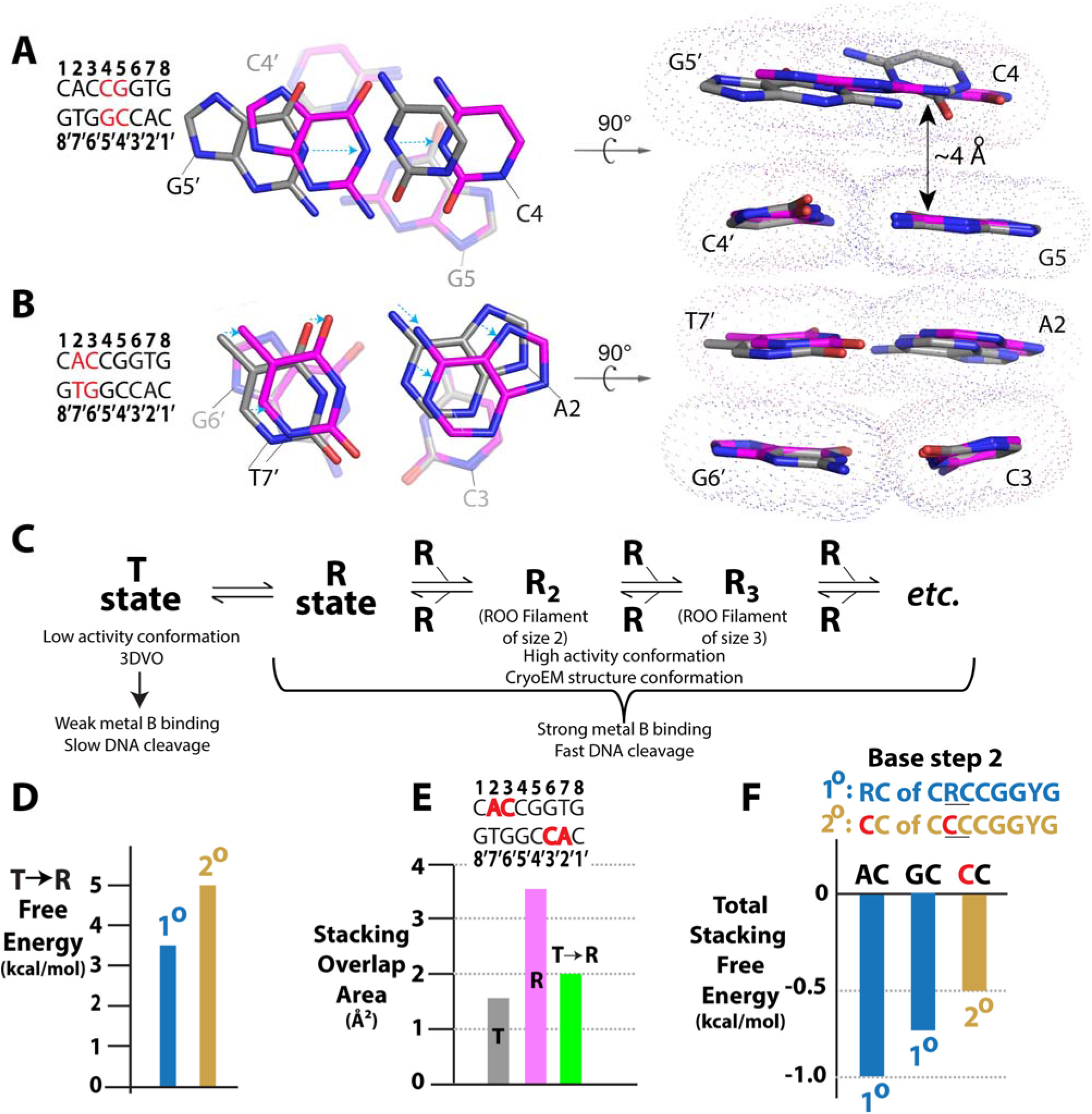
Changes in DNA conformation affect low (T state) and high (R state) conformational energy in DNA sequence dependent manner. **A**. Change in base stacking in T (represented by the non-ROO structure, PDB 3DVO, colored by atom with carbon atoms gray) and the R state (represented by the cryo-EM ROO filament structure, colored by atom with carbon atoms magenta) at the fourth base step. **Left**: Base stack between base pairs C4:G5’ and G5:C4’, highlighted in red in the displayed sequence. Blue dotted arrows indicate shift of base upon the T->R change in conformation. **Right**: View of A rotated by 90°. **B**. As in A, but for the second base step. C. Equilibrium diagram showing the equilibrium between T and R conformational states of SgrAI/DNA complexes. Only the R state has rapid DNA cleavage kinetics. **D**. Estimate of the equilibrium between the T and R states with primary (T state favored by 3.6 kcal/mol) or secondary site (T state favored by 4.9 kcal/mol) bound to SgrAI. **E**. The base stacking (overlap) area (Å^2^) for the T and R states at the second base step, between bases of nucleotides A2 and C3. **F**. Comparison of stacking energies of possible second base step nucleotides. Primary site sequences (blue) provide more stacking energy than secondary (gold)^31^.

To investigate changes in DNA structure further, the stacking overlap areas were calculated for all base steps of the 8-bp recognition sequence (**Table S3**) in both conformations. Similar DNA structure and stacking areas were found at the first and third base steps (**Table S3, Fig. S4A-B**). However, the bases in the second base step, between nucleotides A2 and C3, showed a large rearrangement (**Table S3, Fig. 5B and Movie S2**), corresponding to a change in stacking area of ~2 Å^2^ (a greater than 2-fold difference). This base step is affected by substitutions in primary site sequences (*i.e.* CRCCGGYG) found in two of the fourteen secondary site sequences (*i.e.* C<u>C</u>CCGGYG, Y=C or T). Differences in base stacking are likely to result in differences in stacking energy, and hence different degrees of stabilization of the low and high activity conformations.

### Mechanistic model of SgrAI activation via filament formation and secondary site cleavage activity

One important issue yet to be fully understood is the mechanism by which primary and secondary site sequences influence SgrAI activation. Secondary sites differ from primary sites by a single base pair, either in the first or second position. Some discrimination occurs at the DNA binding step, however SgrAI binds to both types of sites with nanomolar or tigher affinity^10^. Yet cleavage of secondary sites is nearly undetectable unless primary sites are present on the same DNA, or at sufficient concentrations. A working model to rationalize this behavior postulates that SgrAI bound to DNA resides in an equilibrium between a low activity “tense” T state and a high activity “relaxed” R state (**Fig. 5C**). Only the high activity R state forms ROO filaments. When the bound DNA has the primary site sequence, the R state is sufficiently populated to influence ROO filament formation (**Fig. 5C and Fig. 1**). However, when the bound DNA contains the secondary site sequence, the T state is favored to a greater extent, thereby lowering the occupancy of the R state conformation and hence the propensity to form ROO filaments (**Fig. 5C and Fig. 1**). Occupancy of the R state is not completely disallowed, since SgrAI bound to secondary site DNA will join filaments formed by SgrAI bound to primary site DNAs (**Fig. 1**), stabilizing the R state conformation and activating the enzyme for broad cleavage of all bound DNAs.

In the context of the T and R state model, the X-ray crystal structures of non-filamentous SgrAI bound to DNA (primary or secondary site) represent the T state, and the current cryo-EM ROO filament structure of SgrAI bound to primary site DNA represents the R state. We found that the R state is characterized by a reconfiguration of Thr 186 in the enzyme active site, which is predicted to stabilize the site B metal ion (**Fig. 4**). Previous data has shown that SgrAI in the R state exhibits rapid DNA cleavage kinetics, and will form the ROO filament (**Fig. 5C**). The T state has much lower DNA cleavage kinetics, or perhaps, is completely inactive.

The equilibrium between the two states can be estimated by comparing the single turnover DNA cleavage rate constants of SgrAI with primary and secondary site DNAs in the absence of activation and without significant formation of ROO filaments. If only the R state is capable of DNA cleavage (estimated at 0.8 s^-1^)^22^, then the observed cleavage rate constant of the unactivated, non-ROO form of the enzyme bound to primary site DNA (0.0017 s^-1^)^10^ can be used to estimate the proportion of SgrAI/DNA complexes in the T state. This gives an equilibrium constant that shows that the T state is favored over the R state by 470-fold (0.8 s^-1^/0.0017 s^-1^), corresponding to a free energy difference of −3.6 kcal/mol (at 25°C). Since SgrAI cleaves secondary site DNA much more slowly^14^, the T state is favored by this complex to a greater extent, giving −4.9 kcal/mol (**Fig. 5D**). SgrAI bound to either type of site favors the T state conformation, however when the secondary site sequence is bound, the complex favors the T state conformation by an estimated 1.3 kcal/mol more than when the primary site is bound (**Fig. 5D**).

Comparison of the R and T state SgrAI structures bound to primary and to secondary DNA should reveal the origin of the differential stabilization. Only a single structure of SgrAI bound to a secondary site has been determined, that with C<u>C</u>CCGGTG, and in the T state^25^. Previously, no differences in interactions between enzyme and DNA, or in DNA structure, between this and the T state structure with primary site were observed. For this reason, it was necessary to derive a structure of the activated enzyme form in the R state and bound to primary site DNA, which is now represented by our cryo-EM ROO model. The current structure – in particular the configurations of the 1^st^ and 2^nd^ base pairs that account for differences in primary vs. secondary site sequences – was therefore used to investigate possible new interactions or DNA structure that could explain differential activity between primary and secondary site cleavage. The only change in base stacking was identified at the second base step, (C<u>AC</u>CGGTG). Importantly, this subtle change represents a 50% increase in base stacking overlap area in the R state relative to the T (**Fig. 5E**). While the exact energies of base stacking are difficult to calculate, they generally correlate with stacking area. Therefore, this increase in stacking area may be expected to be stabilizing to the R state conformation, relative to the T. Stacking energies measured for different dinucleotide sequences show that the stacking energy of primary sequences at this base step (AC or GC, blue bars, **Fig. 5F**) provide more favorable energy (up to 0.5 kcal/mol) than the secondary site sequence (CC, gold bar, **Fig. 5F**)^31^. Hence, the difference in base stacking and its associated energy may explain at least part of the differential stabilization of the R (vs. T) state by primary and this class of secondary site sequences. A better understanding of the origin in the other class of secondary site sequences (<u>X</u>RCCGGY G, X=G, T, or A) awaits their structure determination in the relevant conformational states.

### Biological role and relationship to other fìlament forming enzymes

The unusual, allosteric, filament forming mechanism of SgrAI may have evolved due to evolutionary pressure imposed by the relatively large genome of its host, *Streptomyces griseus^10^*. The larger genome results in a greater number of recognition sites, which must be protected from SgrAI-mediated cleavage via methylation by the cognate methyltransferase SgrAI.M. Such pressure to protect these sites from damaging cleavage would favor an increase in the activity of the methyltransferase, and/or a decrease in the activity of the SgrAI endonuclease. We find that the activity of SgrAI is in fact reduced compared to that of similar endonucleases, in that its 8-bp recognition sequence is longer than in typical endonucleases, making it rarer in genomes, and its DNA cleavage rate is remarkably slow in the absence of activation^10^. Activation occurs only under particular conditions, namely the assembly of SgrAI into ROO filaments when bound to unmethylated primary site DNA. This is less likely to occur within the host genome, because most primary sites are methylated, but predicted to occur on invading phage DNA containing unmethylated primary sites. In addition to becoming activated upon ROO filament formation, the specificity of SgrAI is also expanded to include cleavage of a second class of recognition sequences, called secondary sites, which increases the total number of recognition sequences from 3 to 17. A greater number of potential cleavage sites increases the probability of phage DNA cleavage, and may also aid in neutralization by preventing transcription, replication, repair, as well as methylation by the SgrAI.M enzyme, also present in the cell.

Beyond the well-known cytoskeletal NTPases, filament formation by enzymes is a newly appreciated phenomenon, with the advantages and evolutionary driving forces yet to be fully understood. Proposed purposes of filament formation vary among enzymes and include rapid enzyme activation, sequestration of activity, buffering of activity, and even in functioning as cytoskeletal structures^5,23,32–35^. In the case of SgrAI, an extensive kinetic study has recently been performed showing that association of SgrAI/DNA complexes into the ROO filament is the rate determining step under most conditions, and is overcome only through high enzyme and DNA concentrations (obtainable *in vitro*). In a biological context, and owing to local concentration effects, filament formation is expected to occur when two cleavage sites reside on the same contiguous DNA^21–23^. This effect is significant and is responsible for sequestering SgrAI activity on phage DNA and away from the host genome^23^. Sequestration of DNA cleavage activity to the activating (*i.e.* phage) DNA is critical, since most secondary sites on the host DNA are not methylated^36^, and can be cleaved by activated SgrAI.

Simulations using the same kinetic parameters determined for SgrAI, but with a theoretical model where only binary oligomeric states are allowed to form, shows that the filament forming mechanism is superior in both speed and sequestration^23^. Both the binary and the ROO filament model effectively sequester activated DNA cleavage on phage DNA, with minimal predicted host DNA cleavage, as a result of the slow, rate-limiting enzyme association into higher-order complexes (*i.e.* the binary complex or ROO filaments). However, the ROO filament mechanism is 2-fold faster in DNA cleavage, owing to the multiple ways enzymes can assemble into filaments (*i.e.* at either end), compared to the binary mechanism with only a single manner of association. The binary mechanism can be “sped up” to achieve the same rate of DNA cleavage as the ROO filament mechanism by increasing the association rate constant for binary assembly formation, however, it must be increased by 4.5-fold to achieve the same fast DNA cleavage kinetics as the ROO filament mechanism^23^. This increased association rate constant results in more predicted DNA cleavage of secondary sites on the host DNA, and hence a loss of sequestration of DNA cleavage activity. Speed is likely to be critical to protecting against phage infection by SgrAI, as is also supported by a recent study^23^. Therefore, the ROO filament mechanism may have evolved to meet the opposing requirements of rapid activation and sequestration of activity.

## METHODS

### Protein preparation

SgrAI enzyme used in assays contains 13 additional C-terminal residues (ENLYFQSHHHHHH) which include 6 histidine residues to be used for SgrAI purification, as well as a cleavage site for TEV protease, and was purified using previously described methods^14^. Briefly, SgrAI was expressed in BL21 (DE3) *E. coli* (which also contain a constitutive expression system for the methyltransferase MspI.M) overnight at 17°C. Cells were sonicated, centrifuged to remove cell debris, and SgrAI was isolated using Talon resin chromatography (Clonetech, Inc.), followed by further purification using heparin resin chromatography (GE, Inc.). Purified SgrAI was concentrated and stored in single use aliquots at −80°C in buffer containing 50% glycerol. Enzyme purity was assessed using Coomassie blue staining of SDS-PAGE and assessed to at least 99% purity.

### DNA preparation

The oligonucleotides were prepared synthetically by a commercial source and purified using C18 reverse phase HPLC. The concentration was measured spectrophotometrically, with an extinction coefficient calculated from standard values for the nucleotides ^37^. Equimolar quantities of complementary DNA were annealed by heating to 90°C for 10 minutes at a concentration of 1 mM, followed by slow cooling to room temperature. The sequence of the DNA used in SgrAI/DNA preparations is shown below (red indicates the SgrAI primary recognition sequence, and | indicates cleavage site):

PC-DNA-top 5’-GATGCGTGGGTCTTCACA −3’

PC-DNA-bottom 3’-CTACGCACCCAGAAGTGTGGCC-5’

Two copies of PC DNA (the duplex formed by annealing of PC-top and PC-bot) self-assemble via annealing of their 5’ “overhanging” CCGG sequences to simulate a 40 bp DNA duplexcontaining a single primary site sequence (shown in red above) after cleavage by SgrAI, however with the exception that it is missing the 5’phosphate at the cleavage site.

### Sample preparation

SgrAI/DNA samples were prepared using: 30 μl of 6.4 μM SgrAI (in 10 mM Tris-HCl pH 7.8, 300 mM NaCl, 0.1 mM EDTA, 1 mM 2-mercaptoethanol), 2 μl of 420 μM stock PC DNA in H_2_O, 1.5 μl of 100 mM Mg(OAc)_2_ and incubation at room temperature for 50 min. The final concentrations are 5.8 μM SgrAI, 25 μM PC DNA (4.3:1 ratio of PC DNA:SgrAI), and 4.5 mM MgCl_2_.

### Data collection

Images were recorded on a Titan Krios electron microscope (FEI) equipped with a K2 summit direct detector (Gatan) at 1.31 Å per pixel in counting mode using the Leginon software package^38^. Data was acquired using a dose of ~55 e^-^/Å^2^ across 60 frames (50 msec per frame) at a dose rate of ~7.8 e^-^/pix/sec. A total of 216 micrographs were recorded over a single session. All imaging parameters are summarized in Table S1.

### Image analysis

Preprocessing steps, including frame alignment, CTF estimation, and particle selection, were performed within the Appion pipeline^39^. Movie frames were aligned using MotionCor2^40^ on 5 by 5 patch squares and using a b-factor of 100. Micrograph CTF estimation was performed using CTFFind4^41^. Particles were selected using DoG picker, with an overlap that did not allow any two picks to be closer than ~150 Å in distance. This provided sufficient separation within the picks to subsequently enable helical averaging of multiple asymmetric units within extracted particle boxes in the 3D classification and refinement stages. 31,988 particles were extracted at this point. Reference-free 2D classification was performed to remove any non-filamentous particles. For 3D classification, initially, an asymmetric (C1) classification was employed to separate helical filaments of distinct compositional heterogeneity and to remove bad particles from the data, which did not give rise to an interpretable reconstruction. At the next step, we imposed helical symmetry during 3D classification to classify the particles. After a round of 2D classification, 3D classification without imposition of helical symmetry, and 3D classification with imposition of helical symmetry, 6,894 particles remained. Finally, 3D refinement was performed using a soft-edged mask, and the resulting map was subjected to B-factor sharpening that yielded a final map of the ROO resolved to ~3.5 Å. The final helical symmetry parameters that were used for refinement were 21.6 Å and −86.2° for the rise and twist, respectively. During refinement, we also determined the optimal number of asymmetric units within a box by iteratively varying the number of asymmetric units (and thus how much helical averaging is performed within each box). Based on examination of the FSC curve and visual inspection of the resulting map, this number was determined to be ~7. Thus, 7-fold helical averaging was performed for each windowed particle. Using this scheme, it is possible that a few of the asymmetric units would appear in the reconstruction more than once, and conversely, that some not at all. However, this was the simplest manner by which to deal with the highly heterogeneous nature of the short, helical fragments that are found in the cryo-EM experiment, while still providing a high-resolution map. The final map was evaluated using Fourier Shell Correlation (FSC) analysis to calculate global and local map resolution (sx_locres.py, implemented within Sparx^42^, provided local FSC maps) and the 3DFSC program suite^43^ to calculate degree of directional resolution anisotropy.

### Atomic model refinement of 3.5 Å resolution cryo-electron microscopy model

**Figure S2** shows the quality of the 3.5 Å cryo-EM envelope. Model building proceeded with real-space refinement after placement of the SgrAI/DNA model derived from X-ray crystallography (PDB 3DVO^18^) using Phenix^44^. Model adjustment and refinement were performed iteratively in Coot^45^ and Phenix^46^, and the statistics were examined using Molprobity^47^ until no further improvements were observed. The final model was also evaluated using FSC analysis against the map (**Figure S1**) and using EMRinger^48^ to compare the fit of the model backbone into the cryo-EM map. The final model statistics showed good geometry and matched the cryo-EM reconstruction (**Figure S1F, Figure S2, and Table S1**).

### Structural comparisons-RMSD and alignments

RMSD calculations were performed with the UCSF Chimera package^49^. Chimera is developed by the Resource for Biocomputing, Visualization, and Informatics at the University of California, San Francisco (supported by NIGMS P41-GM103311). The “Matchmaker” tool (with Needleman-Wunsch matrix for protein, and “nucleic” for DNA) was used with structure-based alignment using alpha carbons, phosphorus atoms, backbone atoms only, or all atoms of selected residues of selected chains. Because the scissile phosphate is missing in the EM structure determined here, the three atoms of the scissile phosphate (the phosphorus atom and two non-esterified oxygen atoms) were deleted from the other structures to allow for superposition with all atoms. PyMOL software (The PyMOL Molecular Graphics System, Version 2.0 Schrödinger, LLC.) was used for figures and some alignments, where indicated, using the align command, or the alignment wizard and selected atoms. Analysis of subunit rotation was performed with UCSF Chimera, after first superimposing both chains A of 3DVO and the cryo-EM model using Matchmaker. The match command was used to calculate the rotation angle of chains B relative to each other.

### DNA conformation and base stacking calculations

The software 3DNA^50,51^ was used to calculate base stacking areas (for all atoms) as well as helical rise.

## Acknowledgments

We thank Yong Zi Tan and Youngmin Jeon for establishing conditions for SgrAI vitrification and Bill Anderson and Jean-Christophe Ducom at The Scripps Research Institute for help with EM data collection and network infrastructure. Molecular graphics and analyses were performed with the USCF Chimera package (supported by NIH P41 GM103331). Research reported in this publication was supported by the National Science Foundation under Grant No. MCB-1410355 (to N.H.) and by the National Institutes of Health Grant No. DP5 OD021396 (to D.L.). The EM map and atomic model of activated filamentous SgrAI will be deposited into the EMDB and PDB under accession codes X and Y respectively.

## Contributions

N.H. prepared and assembled SgrAI and DNA. D.L. collected the cryo-EM data. S.P. processed the cryo-EM data. S.P., N.H., and D.L. built and refined the atomic model; N.H. and D.L. wrote the manuscript; all authors contributed to manuscript editing.

## Supplementary Information

**Figure S1.**
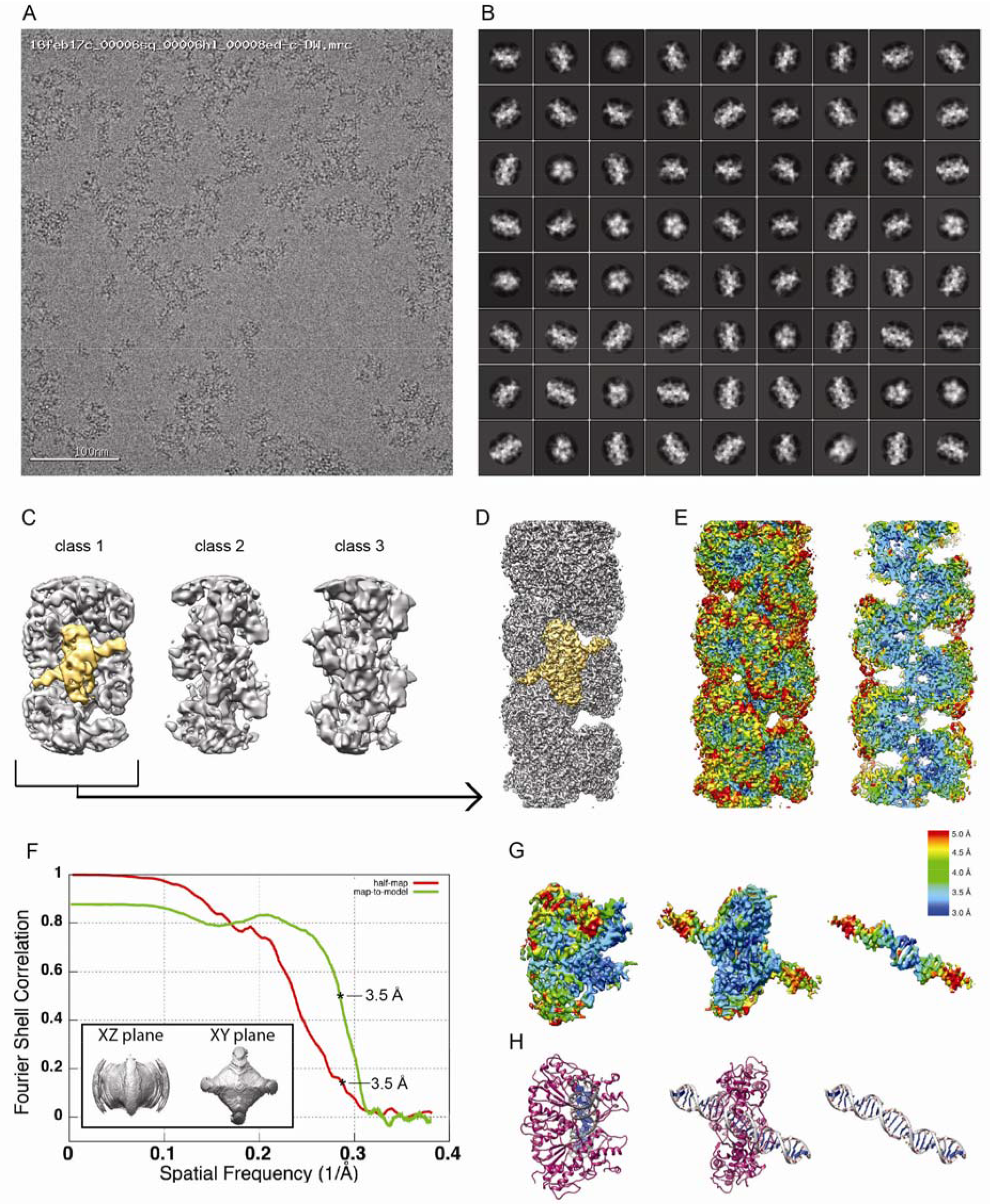
Cryo-EM data. *(A)* Cryo-EM micrograph of SgrAI. *(B)* 2D class averages. *(C)* Reconstructed maps after asymmetric 3D classification showing a class resembling SgrAI (class 1) and two classes with ill-defined features. *(D)* High-resolution refinement of class 1 from C. In both C and D, a single DBD is highlighted in yellow. *(E)* Reconstruction from D colored by local resolution, with the full map at left and a cutaway of the center of the filament at right. *(F)* FSC curves showing half-map resolution (red) and map-to-model resolution (green), both indicating a value of ~3.5 Å. 3D-FSC^43^ isosurfaces are displayed at a threshold of 0.5 within the inset for two perpendicular planar views. *(G)* segmented DBD map colored by local resolution with *(H)* the corresponding atomic model.

**Figure S2.**
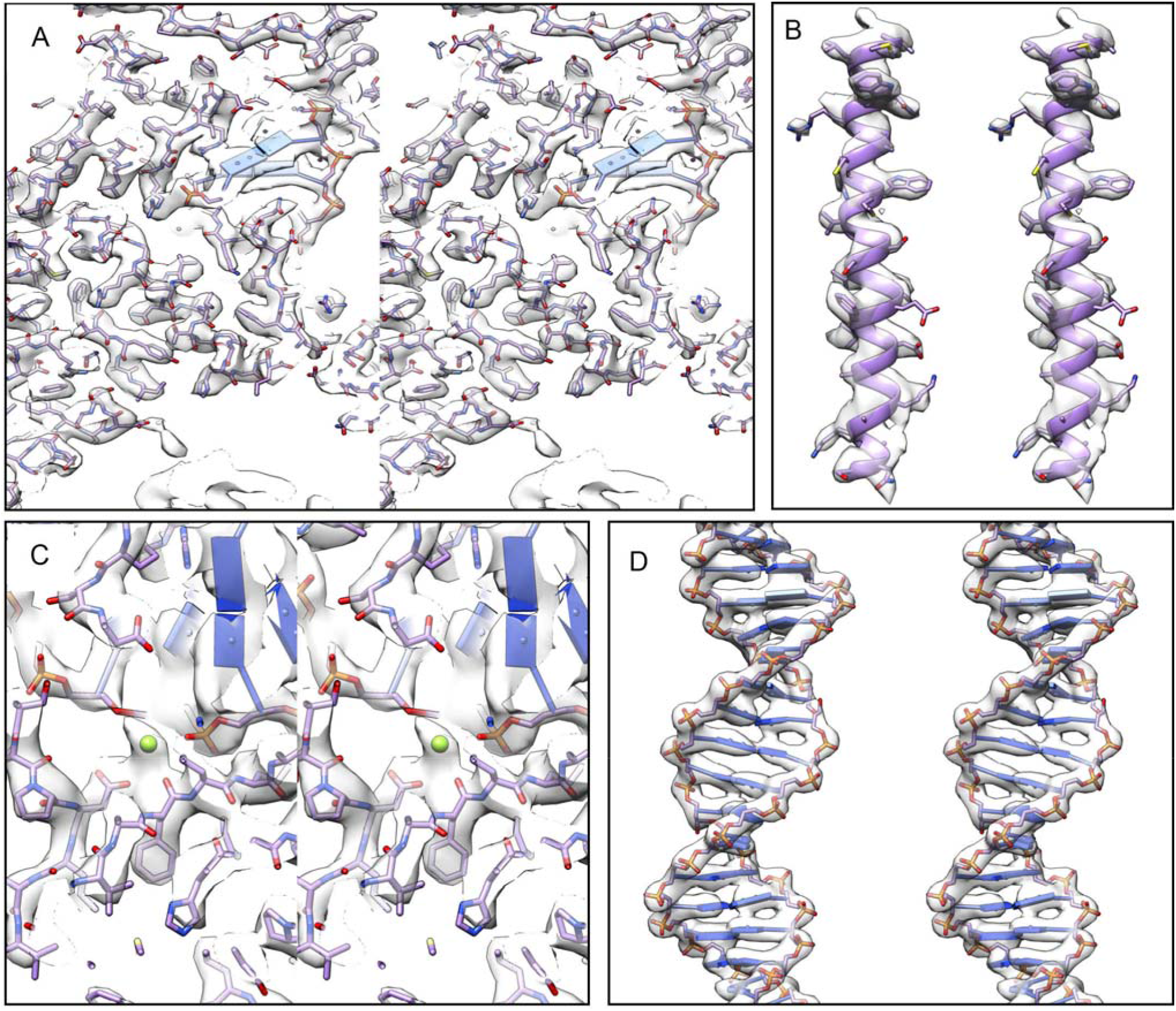
Correspondence between cryo-EM map and atomic model. Stereo views showing *(A)* a section of the ROO, *(B)* an alpha helix (residues 90-122), *(C)* the active site, with bound Mg^2+^ ions (green), and *(D)* segmented density of the PC DNA.

**Figure S3.**
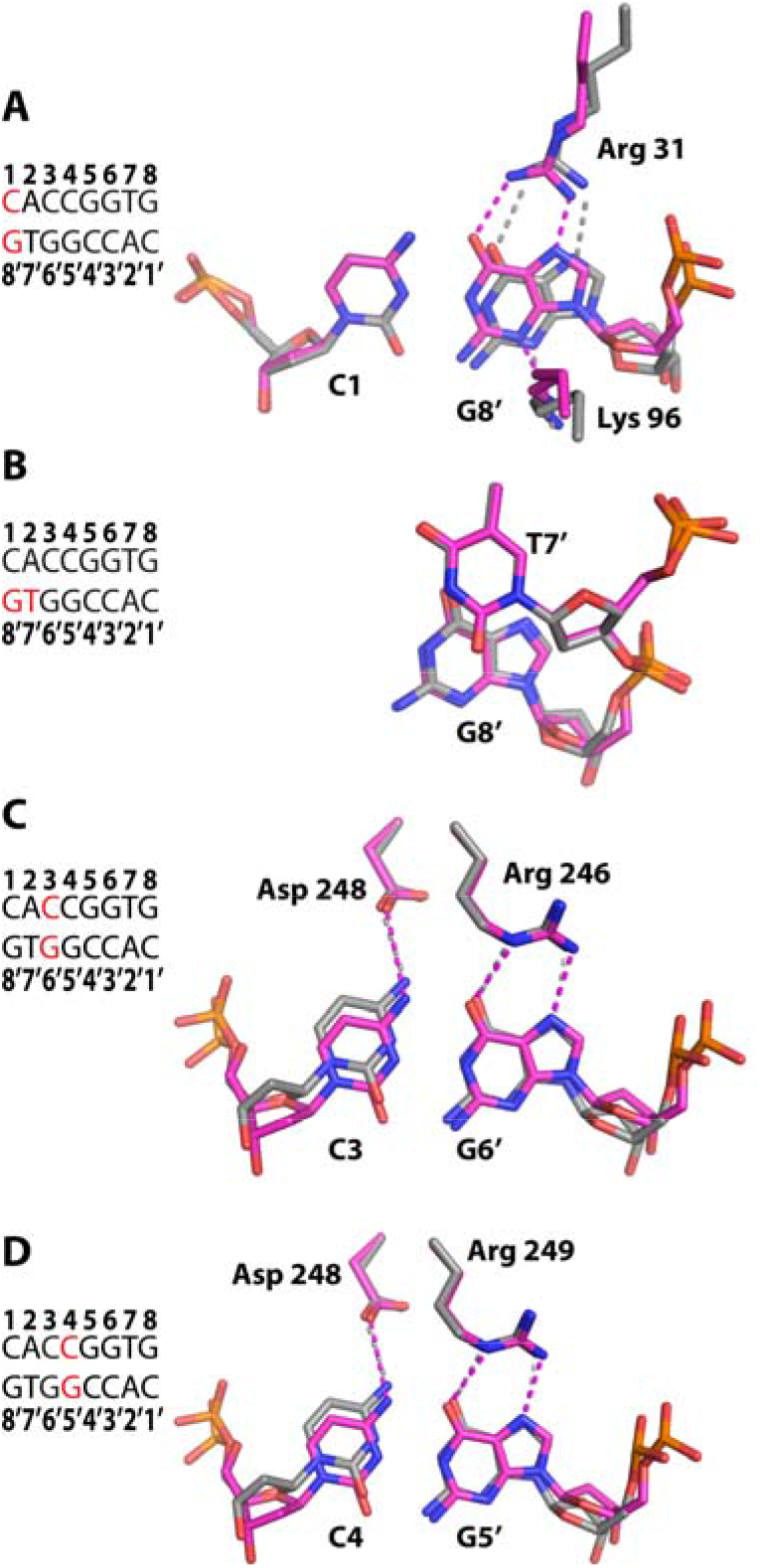
Preservation of contacts between SgrAI and the primary recognition sequence in DNA responsible for sequence specific recognition. *(A)* The low-activity 3DVO structure shown with carbon atoms in gray overlaid with the high-activity ROO filament cryo-EM structure shown with carbon atoms in magenta. Hydrogen bonds indicated by dashed lines. Lys 96 contacts the N3 of G8’, Arg 31 makes hydrogen bonds to the O6 and N7 of G8’. *(B)* Structure at the first base step viewed perpendicular to the base planes. No direct readout contacts are made to the C1 or A2 bases (not shown). Little base stacking (*i.e.* little overlap of the bases in this view) is found between T7’ and G8’, thought to be part of the indirect readout of the pyrimidine at position 7 of the recognition sequence^18^. *(C)* As in A, but contacts to the C3/G6’ base pair. Arg 246 makes hydrogen bonds to the N7 and O6 of G6’. Asp 248 makes a hydrogen bond to the N4 of C3. *(D)* As in A, but contacts to the C4:G5’ bp. Asp 248 makes a hydrogen bond to the N4 of C4, and Arg 249 makes hydrogen bonds to the N7 and O6 of G5’. In all panels, blue, red, and orange refer to nitrogen, oxygen, and phosphorus, respectively.

**Figure S4.**
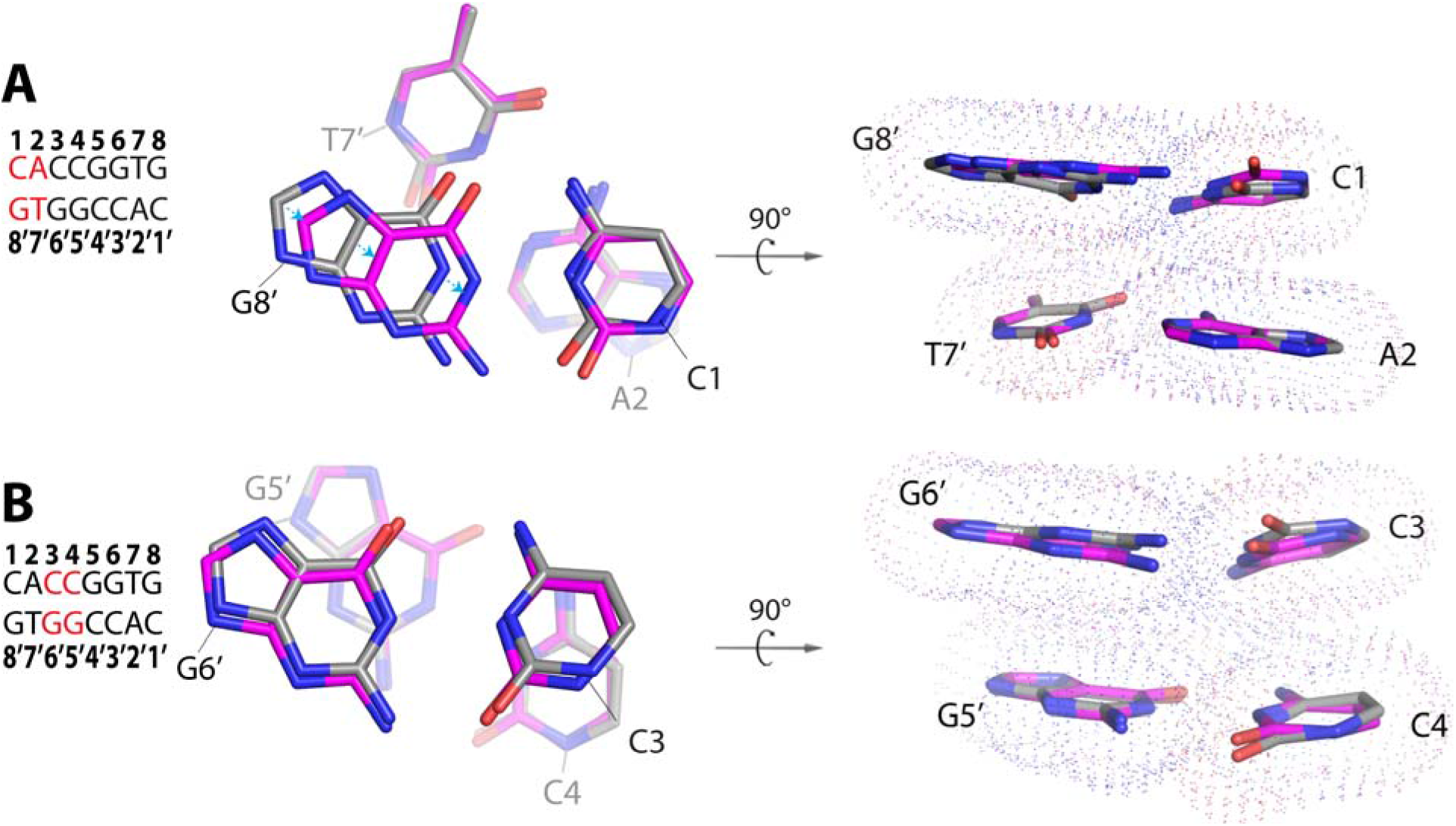
Base stacking comparison between T and R states. *(A)* Base Stacking at the first base step. All atoms of the base pair in the background were used in the superposition. Atoms from the cryo-EM ROO filament model representing the R state shown with carbon atoms in magenta. Those of 3DVO, representing the T state, shown with carbon atoms in grey. A shift in the position of the G8’ base results in a slight increase in stacking overlap with T7’. *(B)* As in A, but the third base step. No significant difference is observed.

**Table S1.**
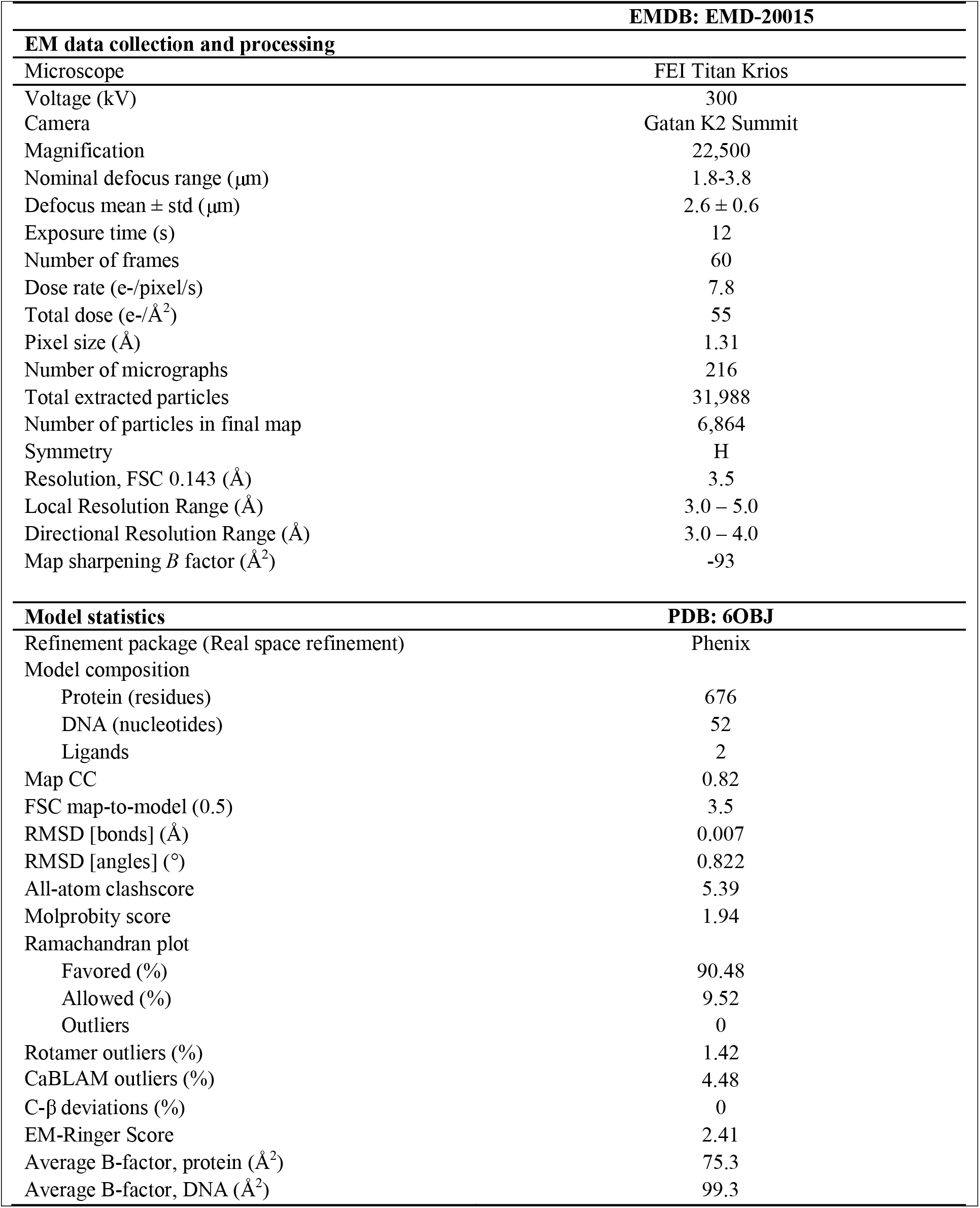
Cryo-EM data collection, refinement, and validation statistics

| **EMDB: EMD-20015** |  |
| --- | --- |
| **EM data collection and processing** |  |
| Microscope | FEI Titan Krios |
| Voltage (kV) | 300 |
| Camera | Gatan K2 Summit |
| Magnification | 22,500 |
| Nominal defocus range ($\mu\text{m}$) | 1.8-3.8 |
| Defocus mean ± std ($\mu\text{m}$) | $2.6 \pm 0.6$ |
| Exposure time (s) | 12 |
| Number of frames | 60 |
| Dose rate (e-/pixel/s) | 7.8 |
| Total dose (e-/$\text{\AA}^2$) | 55 |
| Pixel size ($\text{\AA}$) | 1.31 |
| Number of micrographs | 216 |
| Total extracted particles | 31,988 |
| Number of particles in final map | 6,864 |
| Symmetry | H |
| Resolution, FSC 0.143 ($\text{\AA}$) | 3.5 |
| Local Resolution Range ($\text{\AA}$) | 3.0 – 5.0 |
| Directional Resolution Range ($\text{\AA}$) | 3.0 – 4.0 |
| Map sharpening *B* factor ($\text{\AA}^2$) | -93 |
| **PDB: 6OBJ** |  |
| **Model statistics** |  |
| Refinement package (Real space refinement) | Phenix |
| Model composition |  |
| Protein (residues) | 676 |
| DNA (nucleotides) | 52 |
| Ligands | 2 |
| Map CC | 0.82 |
| FSC map-to-model (0.5) | 3.5 |
| RMSD [bonds] ($\text{\AA}$) | 0.007 |
| RMSD [angles] ($^\circ$) | 0.822 |
| All-atom clashscore | 5.39 |
| Molprobity score | 1.94 |
| Ramachandran plot |  |
| Favored (%) | 90.48 |
| Allowed (%) | 9.52 |
| Outliers | 0 |
| Rotamer outliers (%) | 1.42 |
| CaBLAM outliers (%) | 4.48 |
| C-β deviations (%) | 0 |
| EM-Ringer Score | 2.41 |
| Average B-factor, protein ($\text{\AA}^2$) | 75.3 |
| Average B-factor, DNA ($\text{\AA}^2$) | 99.3 |

**Table S2.** RMSD of DNA between low and high activity conformations of SgrAI/DNA

| Nucleotide of<br>8 bp<br>Recognition<br>Sequence | RMSD<br>(All Atoms,<br>Duplex)<br>(Å<sup>2</sup>) | RMSD<br>(All Atoms,<br>Single<br>Strand)<br>(Å<sup>2</sup>) | RMSD<br>(All Atoms, 4<br>bp Halfsite)<br>(Å<sup>2</sup>) |
| --- | --- | --- | --- |
| C1 | 1.44 | 1.04 | 0.94 |
| A2 | 1.05 | 0.68 | 0.52 |
| C3 | 1.07 | 1.17 | 1.00 |
| C4 | 1.14 | 1.12 | 0.82 |
| G5 | 1.03 | 1.21 | 0.73 |
| G6 | 0.62 | 0.68 | 0.53 |
| T7 | 0.82 | 0.84 | 0.26 |
| G8 | 1.16 | 1.18 | 0.36 |
| All | 1.03 | 0.99 | 0.64 |

**Table S3.** Base Overlap Areas

| Stacked<br>Bases | Area of Low<br>Activity<br>Conformation,<br>T state (3DVO)<br>(Å²) | Area of High<br>Activity<br>Conformation,<br>R state<br>(Cryo-EM ROO<br>Filament) (Å²) | Difference<br>(ROO-3DVO)<br>(Å²) |
| --- | --- | --- | --- |
| **1<sup>st</sup> Base Step**<br>**CACCGGTG**<br>**GTGGCCAC** |  |  |  |
| CA | 5.6 | 5.9 | +0.3 |
| TG | 0.5 | 0.05 | -0.45 |
| **2<sup>nd</sup> Base Step**<br>**CACCGGTG**<br>**GTGGCCAC** |  |  |  |
| AC | 1.6 | 3.6 | **+2.0** |
| GT | 8.2 | 8.6 | +0.4 |
| **3<sup>rd</sup> Base Step**<br>**CACCGGTG**<br>**GTGGCCAC** |  |  |  |
| CC | 6.2 | 6.6 | +0.4 |
| GG | 4.5 | 4.1 | -0.3 |
| **4<sup>th</sup> Base Step**<br>**CACCGGTG**<br>**GTGGCCAC** |  |  |  |
| CG | 2.8 | 1.6 | **-1.2** |
| CG | 2.1 | 1.3 | **-0.8** |
| Cross<br>strand<br>GG | 0 | 0.9 | **+0.9** |

**Movie S1**. Conformational rearrangements between the low- and high-activity SgrAI forms that contribute to helical packing within an ROO

**Movie S2**. Conformational rearrangements between the low- and high-activity SgrAI forms that contribute to an increase in base stacking between the 2^nd^ and 3^rd^ base pairs, and thus sequence specificity expansion.

